# Combined inhibition of apoptosis and necrosis promotes transient neuroprotection of retinal ganglion cells and partial-axon regeneration after optic nerve damage

**DOI:** 10.1101/357566

**Authors:** Maki Kayama, Kumiko Omura, Yusuke Murakami, Edith Reshef, Aristomenis Thanos, Yuki Morizane, Andrea Giani, Toru Nakazawa, Joan W. Miller, Larry Benowitz, Demetrios G. Vavvas

## Abstract

Retinal ganglion cell (RGC) death is the hallmark of glaucoma. Axonal injury is thought to precede RGC loss in glaucoma, and thus studies using an optic nerve (ON) crush model have been widely used to investigate mechanisms of cell death that are common to both conditions. Prior work has focused on the involvement of caspases in RGC death, but little is known about the contribution of other forms of cell death such as necrosis. In this study we show that receptor interacting protein (RIP) kinase-mediated necrosis normally plays a role in RGC death and acts in concert with caspase-dependent apoptosis. The expression of RIP3, a key activator of RIP1 kinase, as well as caspase activity, increased following ON injury. Caspase inhibition alone failed to provide substantial protection to injured RGCs and unexpectedly exacerbated necrosis. In contrast, pharmacologic or genetic inhibition of RIP kinases in combination with caspase blockade delayed both apoptotic and necrotic RGC death, although RGCs still continued to die. Furthermore, inhibition of RIP1 kinase promoted a moderate level of axon regeneration that was only minimal affected by caspase inhibition. In conclusion, multiple approaches are required for effective RGC death prevention and axonal regeneration. Further studies are needed to elucidate more effective long term strategies that can lead to sustained neuroprotection and regeneration.

## Abbreviation List

GCL: retinal ganglion cell layer
IOP: intraocular pressure
IPL: inner plexiform layer
Nec-1: necrostatin-1
ON: optic nerve
PI: propidium iodide
RGC: retinal ganglion cell
RIP: receptor interacting protein
ROS: reactive oxygen species
TEM: transmission electron microscopy
TNF: tumor necrosis factor
WT: wild type

## HIGHLIGHTS

- Caspase inhibition alone does not provide extensive protection against RGC death following optic nerve damage.
- RIP kinase-mediated necrosis is a redundant death pathway in RGCs.
- Combined inhibition of caspases and RIP kinases transiently increases RGC survival.
- Necrosis inhibitors promote a moderate level of axonal regeneration.

## INTRODUCTION

Glaucoma affects 70 million people worldwide (1, 2) and is characterized by optic nerve (ON) atrophy and the progressive death of retinal ganglion cells (RGCs) (3). Glaucoma is often associated with elevated intraocular pressure (IOP), and current management aims at lowering IOP (4). Yet although IOP-lowering treatments slow the development and progression of glaucoma, approximately 10% of people who receive proper treatment continue to experience loss of vision (2). Therefore, understanding the underlying mechanisms involved in RGC death could potentially lead to novel therapeutic approaches.

In general, cell death occurs through two major, morphologically distinct processes, apoptosis, and necrosis (5). Apoptosis is characterized by cellular shrinkage and nuclear condensation, and the caspase family plays a central role in this process (6). In contrast, necrosis is accompanied by cellular and organellar swelling and plasma membrane rupture, and has been considered an uncontrolled form of cell death. However, recent evidence indicates that some necrosis can be executed by regulated signal transduction pathways such as those mediated by receptor interacting protein (RIP) kinases (7, 8). Although previous reports have identified the involvement of apoptosis in RGC death in glaucoma (9), little is known about the contribution of other forms of cell death to the demise of RGCs.

RIP1 is a death domain-containing protein that possesses serine/threonine kinase activity and is a key regulator of cell fate in response to the stimulation of death-domain receptors such as the tumor necrosis factor (TNF) receptor (10). RIP1 forms a death inducing signaling complex with Fas-associated death domain and caspase-8 after stimula-tion of TNF-α, and mediates caspase-dependent apoptosis. On the other hand, when caspases are inhibited or cannot be activated efficiently, RIP1 induces necrosis instead of apoptosis through RIP1-RIP3 binding and phosphorylation of their kinase domain (11–15).

We have previously demonstrated that TNF-α is a significant mediator of RGC death in animal models of glaucoma, with levels rising sharply as a result of elevated intraocular pressure, which leads to an inflammatory response (16) (17). Clinically, TNF-α is up-regulated within RGCs and ON axons of glaucomatous eyes (18), and several other lines of evidence suggest the involvement of TNF-α signaling in glaucoma (19). However, which cell death pathways are utilized in the execution of RGC death are not fully elucidated. Other studies have investigated the role of caspases and their inhibition in protection of RGC cell death and in axonal regeneration (20) (21, 22) (23–25). In general, these studies show that blocking caspases delays but does not fully attenuate the loss of RGCs(26), though a more recent study points to more impressive gains in cell survival (22)

In this study, we investigated the role of RIP-mediated necrosis in addition to the role of caspase-mediated apoptosis and the potential benefit of simultaneous inhibition of these pathways in neuroprotection and axonal regeneration in an ON crush injury model.

## RESULTS

### ON injury induces caspase activation and increases RIP kinase expression

The initiation of RGC death after ON damage has been linked to the disruption of retrograde axonal transport (27) and activation of the dual-leucine kinase pathway (28, 29), and is widely used as a model of RGC death in glaucoma (30). Here, we evaluated changes in known effectors of cell death following optic nerve injury, including caspases as mediators of apoptosis and RIP kinase as a mediator of necrosis (10). One day after ON injury (vehicle), the activities of caspase-8, -9 and -3 were significantly increased compared with those in non-injured retina (Figure 1A). RIP3 is a key regulator of RIP1 kinase activation, and its expression level has been shown to correlate with responsiveness to programmed necrosis (12). Quantitative real-time PCR analysis revealed that RIP3 and RIP1 expression in the retina increased up to 9- and 5-fold, respectively, 1 day after ON injury compared with those in non-injured retina (Figure 1B). Western blot analysis of retina extracts confirmed the up-regulation of RIP3 and RIP1 expression after ON injury compared with non-injured retina (Figure 1C). These results suggest that both caspase and RIP kinase pathways may contribute to RGC death after ON injury.

**Figure 1.**
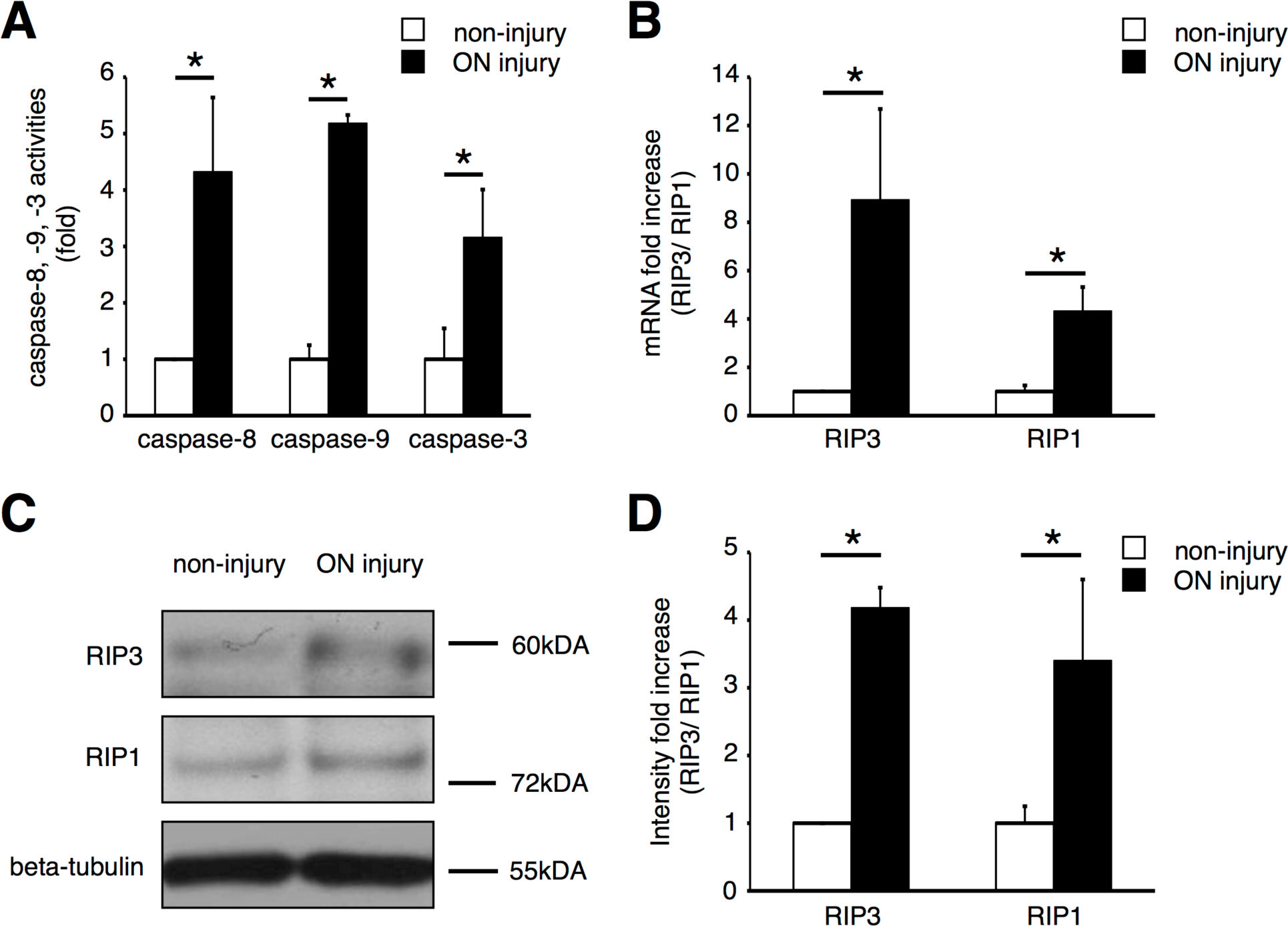
Changes in caspase and RIP kinase pathways after ON injury. **A.** Caspase-8, -9 or -3 activities in the retina after ON injury. These activities were up-regulated 1 day after ON injury (*n* = 6, \**p* < 0.01). The caspase activities were normalized by non-injured retina. **B.** Quantitative real-time PCR analysis of RIP3 and RIP1 expression in the retina after ON injury. Expressions of RIP3 and RIP1 significantly increased 1 day after ON injury (black box) compared with those in non-injured retina (white box) (*n* = 6, \**p* < 0.01). **C and D**, Western blot analysis for RIP3 and RIP1 **(C)**. Relative protein expression ratio of RIP3 and RIP1 at 1 day after ON injury **(D)**. The protein expression levels of RIP3 and RIP1 were normalized by non-injury. RIP3 and RIP1 expressions were increased after ON injury (*n* = 6, \**p* < 0.01). Lane-loading differences were normalized by levels of ®-tubulin.

### Individual and combined treatment of caspase and RIP1 kinase inhibitor suppresses RGC death after ON injury

To further investigate the role of caspases and RIP kinases in RGC death after ON injury, we used a pan-caspase inhibitor, ZVAD (benzoyl-Val-Ala-Asp-fluoromethyl ketone) and a RIP1 kinase inhibitor, Necrostatin-1 (Nec-1). TUNEL (+) cells in the retinal ganglion cell layer (GCL) were assessed 1 day after ON injury, because the number of TUNEL (+) cells peaked at this time point (Figure 2A and 2B). Although TUNEL has been used as a marker of apoptosis, several studies have indicated that necrosis, programmed or otherwise, also yields DNA fragments that react with TUNEL, rendering it difficult to distinguish between apoptosis and necrosis (14, 31). Intravitreal injection of ZVAD (300 μM) decreased the activities of caspase-8, -9 and -3 by approximately 60%; however, caspase inhibition with ZVAD did not significantly alter the number of TUNEL (+) cells (Figure 2A and 2B). Administration of Nec-1 alone also failed to show a significant protective effect. However, co-treatment with ZVAD and Nec-1 resulted in a dramatic decrease of TUNEL (+) cells 1 day after ON injury (Figure 2A and 2B).

**Figure 2.**
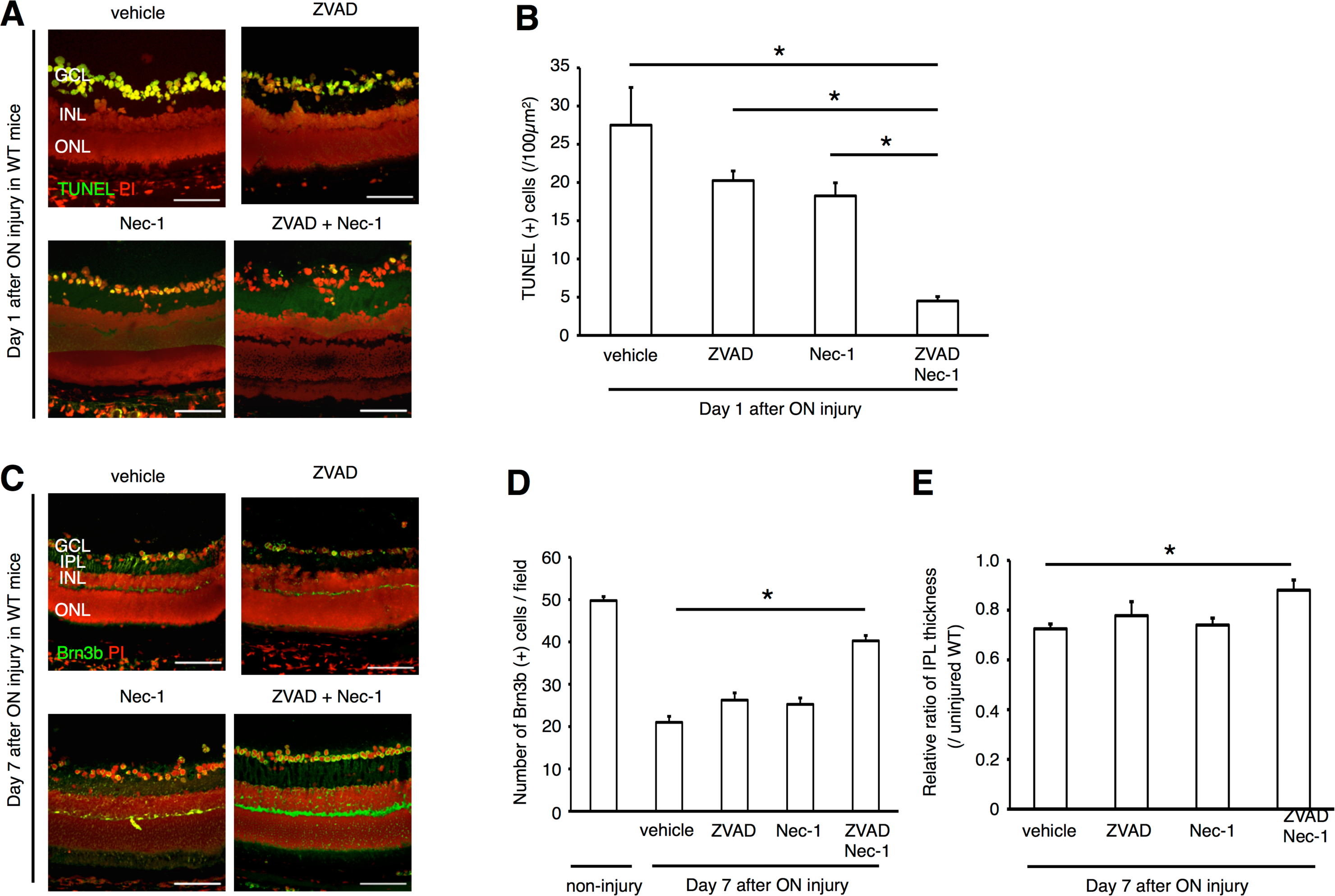
Co-administration of ZVAD plus Nec-1 prevents RGCs death after ON injury. **A and B**, TUNEL assay **(A)** and quantification of TUNEL (+) cells **(B)**. The eyes treated with vehicle, ZVAD, Nec-1 or ZVAD plus Nec-1 were analyzed 1 day after ON injury (*n* = 4). The number of TUNEL (+) cells in GCL was significantly decreased by treatment of ZVAD plus Nec-1 (\**p* < 0.01). **C-E**, Immunofluorescent staining for Brn3b (green) and PI (red) **(C)**, quantification of Brn3b (+) cells **(D)** and IPL thickness ratio **(E)**. The decrease of Brn3b (+) cells and IPL thickness ratio after ON injury was markedly suppressed by treatment of ZVAD plus Nec-1 (*n* = 6–8, *\*p* < 0.01). Bars: **(A, C)** 100 ⌠ m. GCL; retinal ganglion cell layer, IPL; inner plexiform layer, INL; inner nuclear layer, ONL; outer nuclear layer.

We next evaluated the injury and loss of RGCs after ON damage by performing immunofluorescence for the specific RGC marker, Brn3b, which is downregulated after nerve damage and precedes RGC loss, and by measuring inner plexiform layer (IPL) thickness. Significant decreases of Brn3b (+) expression and IPL thickness ratio were first observed at 3 days after ON injury and progressed by 7 days (Figure 2C-2E) Treatment with ZVAD or Nec-1 alone failed to rescue RGC loss at 7 days after ON injury, However, co-administration of ZVAD and Nec-1 markedly suppressed the decrease in Brnb3 (+) cells and IPL thickness ratio (Figure 2C-2E). Administration of ZVAD alone, Nec-1 alone, or ZVAD plus Nec-1 did not affect the TNF-α expression levels after ON injury. Taken together, these data suggest that inhibition of both caspases and RIP1 kinase is required for effective neuroprotection after ON injury, at least in the short term.

### Programmed Necrosis is involved in RGC death after ON injury and is exacerbated by inhibition of apoptosis

Next, we investigated the morphology of dying RGCs by transmission electron microscopy (TEM). RGC death was categorized into apoptosis, necrosis and unclassified end-stage of death, as previously described (14). One day after ON injury, both apoptotic and necrotic RGCs were observed in the vehicle-treated retina (apoptotic cells: 13.4 ± 5.8%, necrotic cells: 16.9 ± 4.2%, unclassified: 2.2 ± 2.4%; Figure 3A and 3B). Nec-1 treatment slightly decreased necrotic RGC death (% apoptotic cells: 13.0 ± 8.4%, necrotic cells: 10.6 ± 2.6%, unclassified: 1.0 ± 1.8%; Figure 3A and 3B). In contrast, ZVAD treatment significantly decreased apoptotic RGC death, while it increased necrotic cell death without reducing overall cell loss (% apoptotic cells: 5.6 ± 3.4%, *P* < 0.01; necrotic cells: 30.7 ± 4.9%, *P* < 0.01; unclassified: 3.4 ± 2.3; Figure 3A and 3B). In addition, infiltration of inflammatory cells was more prevalent in ZVAD-treated retina (Figure 3A). Co-administration of ZVAD and Nec-1 led to a substantial decrease of both apoptotic and necrotic RGC death (% apoptotic cells: 4.2 ± 5.1%, *P* < 0.01; necrotic cells: 9.8 ± 6.6%, *P* < 0.01; unclassified: 1.8 ± 1.8%; Figure 3A and 3B).

**Figure 3.**
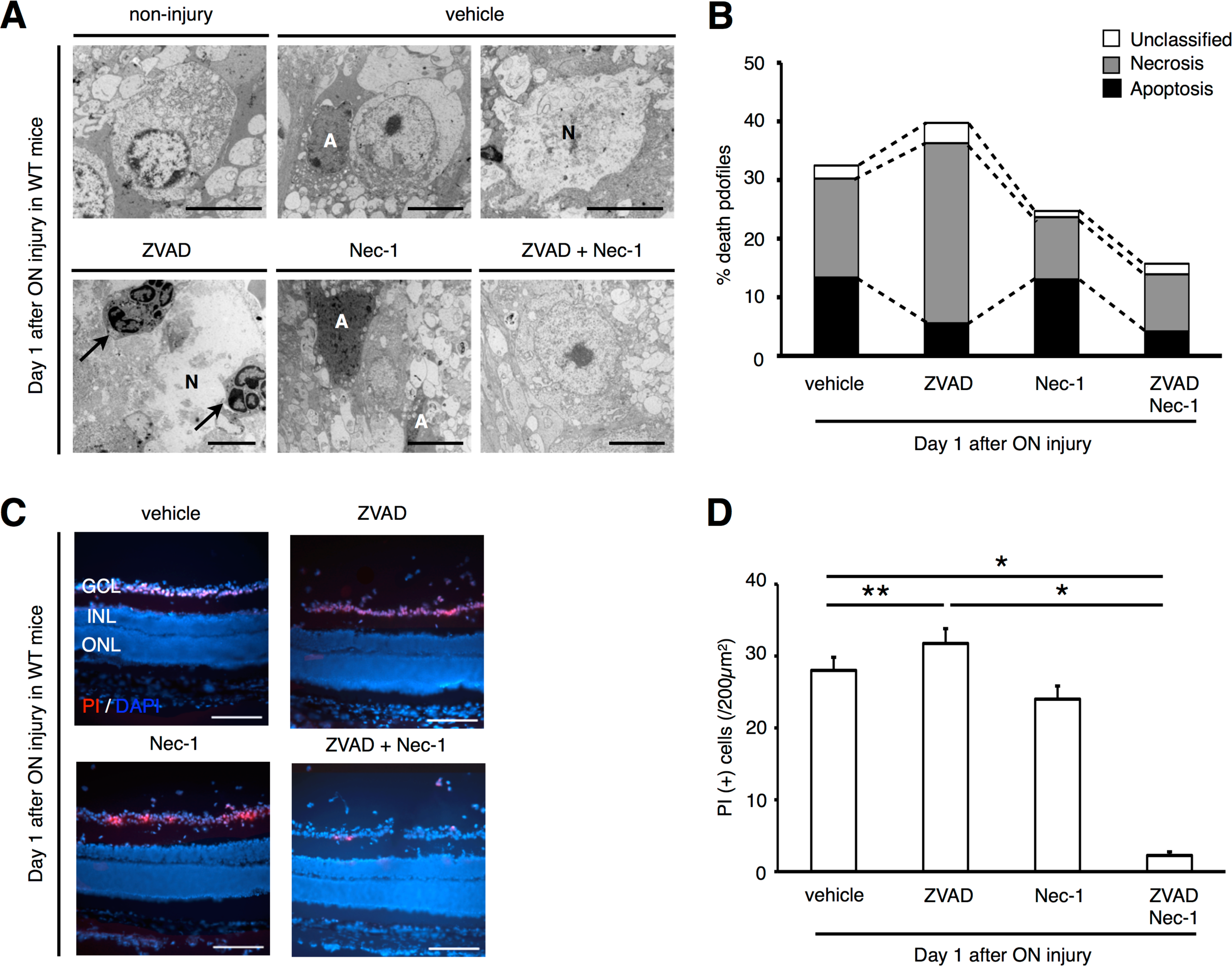
Programmed necrosis is involved in RGC death after ON injury and compensates for inhibition of apoptosis. **A and B**, TEM photographs of RGCs 1 day after ON injury **(A)** and quantification of apoptotic and necrotic RGC death **(B)**. The eyes were treated with vehicle, ZVAD, Nec-1 or ZVAD plus Nec-1. Both apoptotic and necrotic cell death was observed after ON injury (vehicle). ZVAD-treated retina showed increased necrosis and infiltration of inflammatory cells (arrows). Co-administration of ZVAD plus Nec-1 led to reduction of both apoptotic and necrotic RGCs. A: Apoptosis, N: Necrosis. **C and D**, *In vivo* PI staining (red) (C) and quantification of PI (+) cells **(D)** in the GCL by immunohistochemistry. DAPI (blue) was used for nuclear staining. Note that the number of PI (+) cells was significantly increased in ZVAD-treated group than in vehicle-treated group. Co-administration of ZVAD plus Nec-1 significantly reduced the number of PI (+) cells (*n* = 6, \**p* < 0.01, \*\**p* < 0.05). Bars: **(A)** 5⌠ m, **(C)** 200⌠ m. GCL; retinal ganglion cell layer, INL; inner nuclear layer, ONL; outer nuclear layer.

Consistent with TEM results, intravitreal injection of propidium iodide (PI), a membrane-impermeant dye which detects cells with disrupted cell membrane (32), showed increased numbers of PI (+) cells in ZVAD-treated retina compared with vehicle-treated retina after ON injury (Figure 3C and 3D). In contrast, the number of PI (+) cells was significantly decreased by co-administration of ZVAD plus Nec-1 (Figure 3C and 3D). Collectively, these results demonstrate that RIP1 kinase-mediated necrosis is an important pathway of RGC death in addition to apoptosis, and compensates for blockade of caspase-dependent apoptosis after ON injury.

### *Rip3* deficiency prevents necrosis induction and attenuates RGC loss after ON injury

To further elucidate the role of RIP1 kinase in RGC death after ON injury, we used mice deficient for *Rip3*, since *Rip1*^*-/-*^ mice die postnatally at day 1-3 (33). *Rip*3^*-/-*^ mice exhibited significantly fewer TUNEL (+) cells 1 day after ON injury compared with wild type (WT) controls (Figure 4A and 4B). ZVAD administration in *Rip3*^*-/-*^ mice led to a further decrease in TUNEL (+) cells compared with vehicle-treatment, whereas Nec-1 administration did not have any additional effect (Figure 4A and 4B). The decrease in Brn3b (+) cells and IPL thickness 7 days after ON injury was significantly suppressed in *Rip3*^-/-^ mice, and this protective effect was further enhanced by ZVAD administration (Figure 4C-4E). In contrast, administration of Nec-1 alone in *Rip3*^*-/-*^ mice did not show any additional effect.

**Figure 4.**
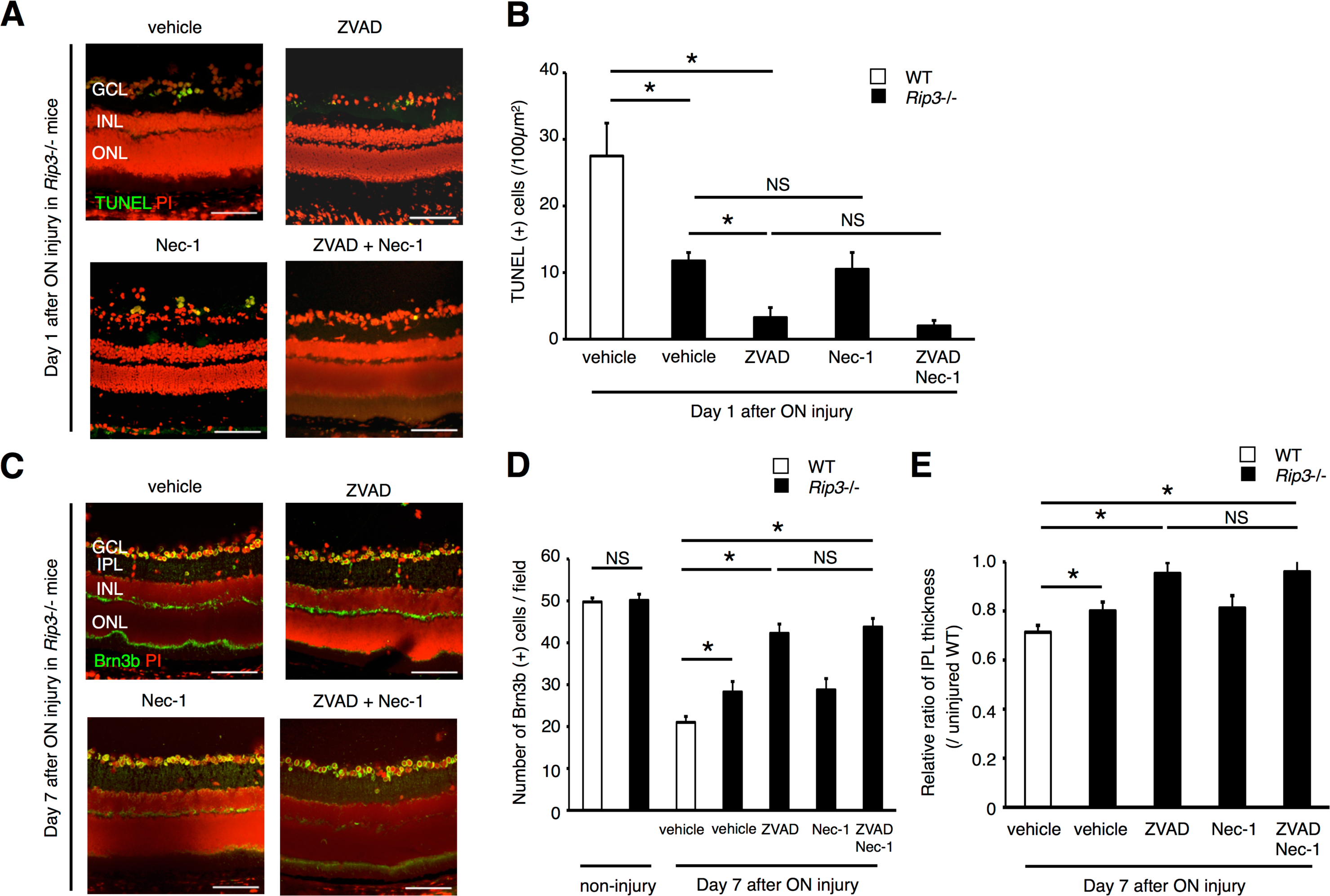
*Rip3* deficiency attenuates RGC loss after ON injury. **A and B**, TUNEL assay **(A)** and quantification of TUNEL (+) cells in WT (white box) and *Rip*3^*-/-*^ mice (black box) **(B)**. The eyes treated with vehicle, ZVAD, Nec-1, or ZVAD plus Nec-1 were analyzed 1 day after ON injury. *Rip*3^*-/-*^ mice exhibited significantly less TUNLE (+) cells than WT mice, and this protective effect was augmented by ZVAD treatment (*n* = 4, \**p* < 0.01, NS; not significant). **C-E**, Immunofluorescent analysis for Brn3b (green) and PI (red) **(C)**, quantification of Brn3b (+) cells **(D)** and IPL thickness ratio **(E)**. The decrease of Brn3b (+) cells and IPL thickness ratio after ON injury was suppressed in *Rip*3^*-/-*^ mice compared with that in WT mice. ZVAD treatment in *Rip*^*-/-*^ mice further prevented the reduction of Brn3b (+) cells and IPL thickness ratio (*n* = 4, \**p* < 0.01, NS; not significant). Bars: **(A, C)** 100⌠ m. GCL; retinal ganglion cell layer, IPL; inner plexiform layer, INL; inner nuclear layer, ONL; outer nuclear layer.

These data confirm that RIP kinases play an essential role in the induction of necrosis after ON injury.

### RIP kinases mediate ROS production after ON injury

Recent studies have shown that RIP kinases induce programmed necrosis via reactive oxygen species (ROS) overproduction (11, 13, 34). One day after ON injury, retinal oxidative damage, as assessed by ELISA for carbonyl adducts of proteins, increased significantly compared with untreated retina (Figure 5). Interestingly, ZVAD administration enhanced retinal oxidative damage, suggesting that RIP kinase activation may promote ROS overproduction (14). Indeed, co-administration of ZVAD and Nec-1 or *Rip3* deficiency substantially decreased retinal oxidative damage after ON injury (Figure 5). These results indicate that RIP kinases play a crucial role in ROS production.

**Figure 5.**
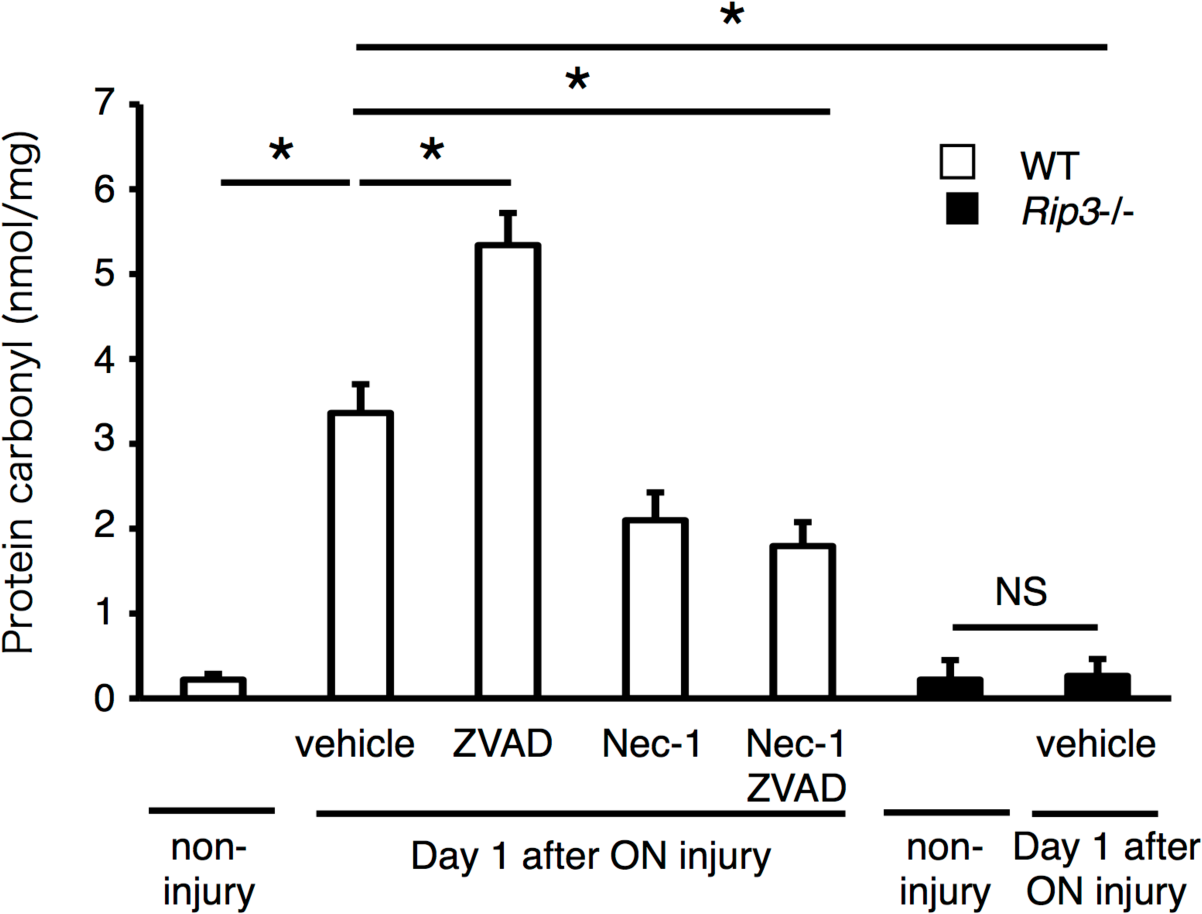
RIP kinase inhibition prevents ROS production after ON injury. ELISA to evaluate carbonyl contents 1day after ON injury. Non-injured retina was used as a control (*n* = 4, \**p* < 0.01, NS; not significant). WT (white box) or *Rip3*^*-/-*^ mice (black box) were treated with vehicle, ZVAD, Nec-1 or ZVAD plus Nec-1, and the retinas were collected 1 day after ON injury. After ON injury, carbonyl contents increased, and this increase was further enhanced by ZVAD. Nec-1 or *Rip*3 deficiency significantly reduced the increase of carbonyl contents after ON injury (*n* = 4, \**p* < 0.01, NS; not significant).

**Fig 6.**
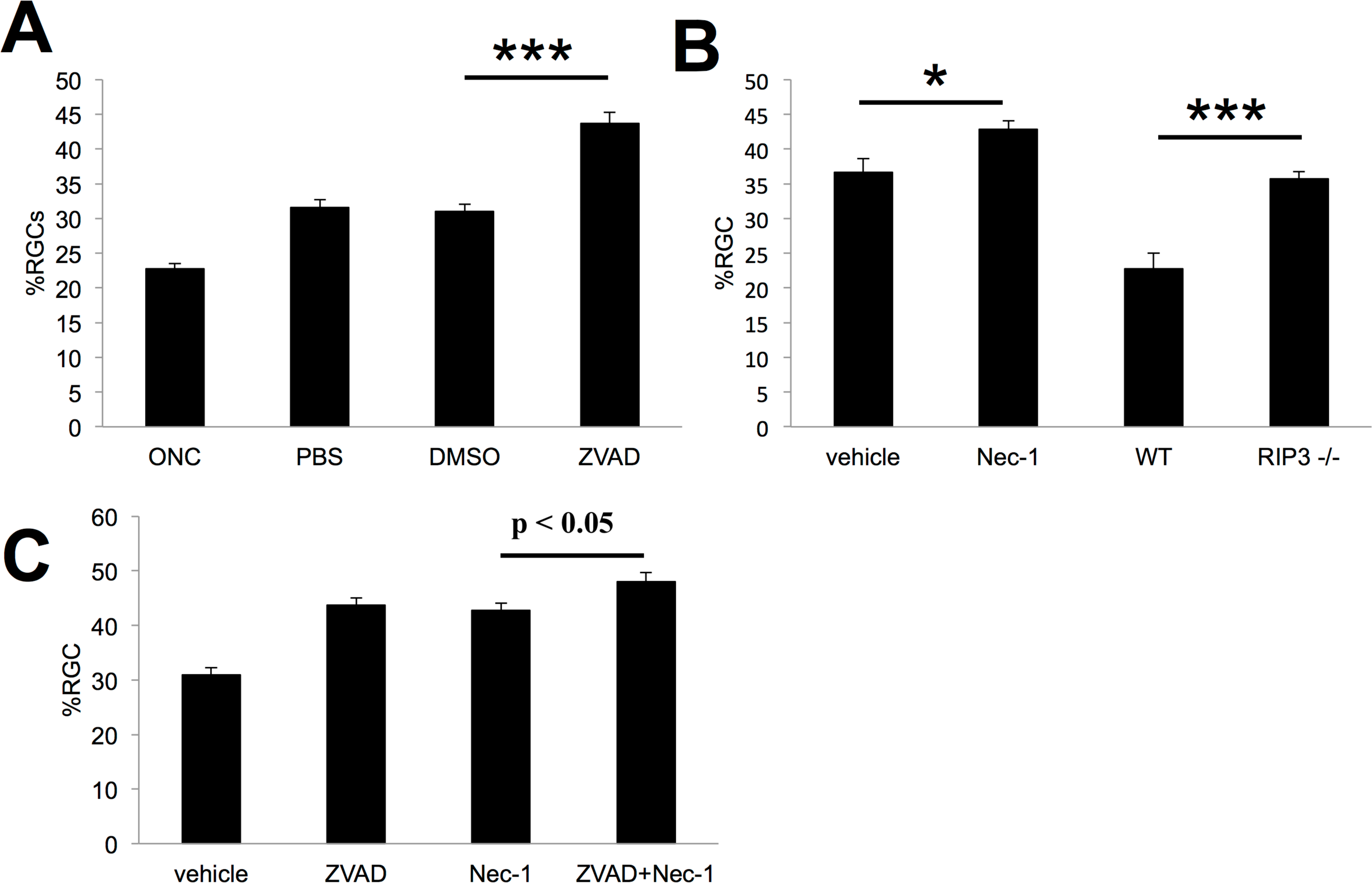
Long term RGC survival after Caspase and RIPK inhibition. Caspase Inhibitors (zVAD) and RIP1K inhibitor (Nec1) were administered intravitreally immediately after nerve injury and again after 3 days and 7 days. RGC survival was examined at 2 weeks post-crush. Repeated injections of vehicle alone increased RGC survival. Repeated intraocular injections of ZVAD increased survival over 40% (Fig. 6A:*P* < 0.001). Repeated injections of the vehicle used for Nec-1 itself increased survival, and the addition of Nec-1 only augmented survival by another 6% (Fig. 6B:*P* < 0.05). Similarly, *Rip3*^*-/-*^ mice exhibited an increase in RGC survival relative to untreated wild-type mice, bringing overall survival only up to 35% normal. (Fig. 6B:*P* < 0.001). In addition, in contrast to our short-term experiment, the combination of Nec-1 and ZVAD caused only a marginal increase in RGC survival relative to either Nec-1 or ZVAD alone ((Fig. 6C *P*= 0.08). (n=6–8 for each group)

**Fig 7.**
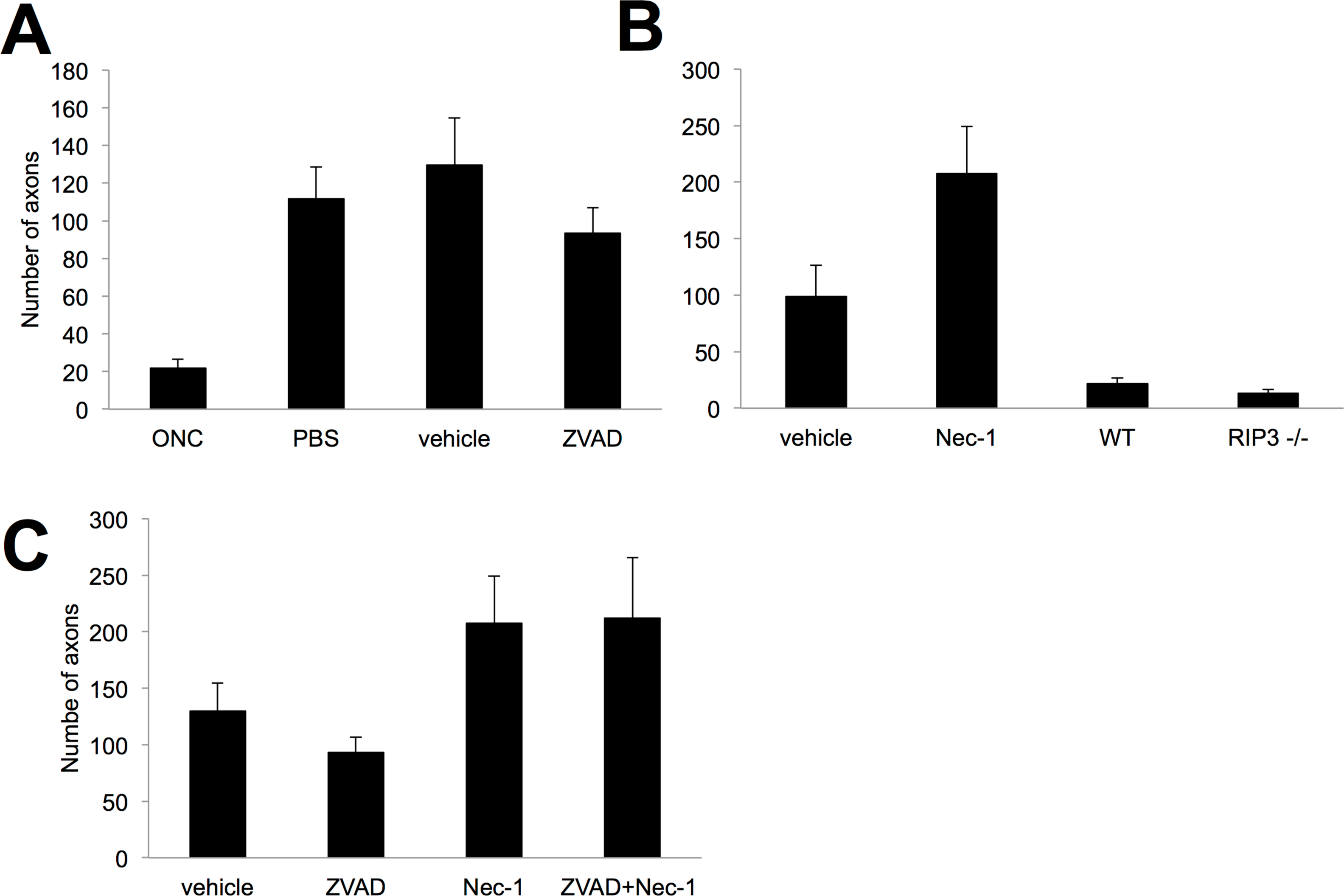
Necrostatin increases Axonal Regeneration independently of Rip3 inhibition. The general caspase inhibitor ZVAD caused no additional increase in regeneration over the vehicle control (7A). In contrast, the RIPK1 inhibitor Nec-1 led to more axon regeneration than the vehicle control (7B). The addition of ZVAD to Nec-1 did not enhance this effect (Fig. 7C). To investigate whether the effects of Nec-1 were mediated through the RIP1-RIP3 interaction, we examined regeneration in *Rip*^*-/-*^ mice. To our surprise the RIP3 null mice did not exhibit any axonal regeneration, suggesting that other effectors of RIP1K or other targets of Nec-1 may mediate the effects of that agent on axonal regeneration (Fig. 7B). (n=6–8 for each group)

### Long-term effects of inhibiting Caspases and RIP kinases after ON injury

Given the increased effectiveness of combinatorial treatment noted in the short term (1-7 days), we investigated whether combinatorial treatment also provides longer-term protection for RGCs. For these studies, we applied agents at three time points (immediately after nerve injury and again after 3 days and 7 days) and examined RGC survival at 2 weeks post-crush. As seen in Fig 6A repeated injections of vehicle alone increased RGC survival by 40% over baseline (Normal RGC baseline numbers: 2895/sq.mm), and repeated intraocular injections of ZVAD increased survival another 40% above this level (Fig. 6A, *P* < 0.001). Repeated injections of the vehicle used for Nec-1 alone increased survival by 60% above baseline, and the addition of Nec-1 only augmented survival by another 17% to a total of 44% of normal. Similarly, *Rip3*^*-/-*^ mice exhibited a 64% increase in RGC survival relative to untreated wild-type mice, bringing overall survival only up to 36% normal. Because these mice lack the *rip3k* gene entirely, the extensive cell loss seen at 2 weeks implies that there is only a minor long-term benefit of preventing necrotic cell death. In addition, in contrast to our short-term experiment, the combination of Nec-1 and ZVAD caused only a marginal increase in RGC survival relative to either Nec-1 or ZVAD alone in the long term (P = 0.08). This result may be due to rapid clearance of these small molecules, or that long-term survival requires suppression of additional cell-death pathways.

### Necrostatin aids axonal regeneration

Inhibitors of caspases-6 and -8 were recently reported to increase the sprouting of injured axons in the optic nerve (20), and even stronger effects were reported after inhibiting Caspases-2 and -6(22). In our hands, repeated injections of either saline or 1% DMSO were sufficient to induce considerable regeneration, probably due to the induction of an inflammatory response in the eye and subsequent elevation of the atypical growth factor oncomodulin (35) (36, 37). The general caspase inhibitor ZVAD caused no additional increase in regeneration (Fig. 7A) nor did repeated injections of either z-IETD-FMK, a specific inhibitor of caspase-8, or z-DEVD-FMK, a specific inhibitor or caspase-3 (Suppl. Fig 1). In contrast, the RIPK1 inhibitor Nec-1 led to substantially more axon regeneration than the vehicle control (Fig 7B). The addition of ZVAD to Nec-1 minimally enhanced this effect (Fig. 7C).

To investigate whether the effects of Nec-1 were mediated through the RIP1-RIP3 interaction, we examined regeneration in *Rip3*^*-/-*^ mice. To our surprise the RIP3 null mice did not exhibit any axonal regeneration, suggesting that other effectors of RIP1K or other targets of Nec-1 may mediate the effects of that agent on axonal regeneration (Fig. 7B).

## DISCUSSION

Considerable evidence suggests that in glaucoma, damage begins within the ON due to structural changes within the lamina cribrosa (38), leading to cellular changes that influence RGC viability (39). Thus axonal injury has long been studied as model of glaucoma. Accumulating evidence suggests that TNF-α contributes to neuronal death in glaucoma (19). TNF-α is elevated in the aqueous humor of glaucoma patients (40), and polymorphisms in TNF-α - related genes are associated with increased risk for glaucoma (41). Neutralization of TNF-α or deleting the gene encoding TNF-α or one of its receptors (TNFR2) prevents RGC death in several experimental models of glaucoma (16, 19) (17). However, the mechanism by which TNF-α signaling leads to RGC death remains unclear, and may involve inflammation and Fas ligand as intermediaries(16, 17, 42-48). Other studies have investigated the effects of inhibiting caspases in protecting RGCs and promoting axonal regeneration (20) (21) (22). In general, these studies show that blocking caspases delays but does not fully attenuate the loss of RGCs, and may have some effects on axonal regeneration (20) (21) (22). Over the past years there has been increasing awareness of caspase-independent death pathways and appreciation that cells can die though alternate pathways such as RIPK mediated necrosis. (7, 8, 49) (11–14), (15). In our study, we found that expression of RIP1 and RIP3, a key activator of RIP1 kinase, was significantly elevated after ON Crush. Tezel *et al* reported an up-regulation of RIP kinase expression in primary RGC culture after 24hr exposure to TNF-α using cDNA arrays (50). Furthermore, we demonstrated that Nec-1, which targets RIP1 phosphorylation, and Rip3 deficiency (11, 51), suppressed RGC death when combined with pan-caspase inhibitor. However, at later time points, RGCs continued to die, even when Nec-1 was combined with caspase inhibitors. Thus, although RIP1 kinase contributes to RGC death, alternative death pathways that are unaffected by our treatments appear to remain active.

Although the presence of necrosis as well as apoptosis was reported in previous morphological analysis of RGC death in experimental glaucoma (52, 53), most recent studies have not focused on necrosis since it has been considered a passive, unregulated form of cell death. In this study, we showed that blockade of caspases with ZVAD decreased apoptosis but exacerbated necrotic RGC death. Furthermore, early necrotic changes were reversed after Nec-1 co-administration. These findings clearly identify RIP kinase-mediated necrosis as an important and complementary mechanism of RGC death that acts in parallel with caspase-dependent apoptosis.

Most previous studies on neuroprotection have shown the effect to be short term. We hypothesized that combination therapy might result in longer lasting effects. However our experiments demonstrated that our long-term effects were muted. This could be a result of pharmacokinetics, since both caspase inhibitors and necrostatins are small molecules with short half-life (in hours), which precludes frequent injection into the mice eyes. Morever, systemic administration is only feasible for Necrostatin and not the combination, since zVAD does not cross the Blood Brain Barrier. In addition our data suggest that there may be more pathways involved in long term survival of RGCs. An important downstream event of RIP kinase activation is the generation of ROS, which is thought to contribute to necrotic and autophagic cell death. RIP1 kinase has been reported to regulate ROS production both directly and indirectly. First, after TNF-α stimulation, RIP1 forms a complex with NADPH oxidase 1 and produces superoxide during necrosis (34). Second, activated RIP3 interacts with metabolic enzymes such as glutamate dehydrogenesel and thereby increases mitochondrial ROS production (13). Third, RIP1 kinase activates autophagic degradation of catalase leading to ROS accumulation (54). In fact, we observed that ROS accumulation after ON injury increased after ZVAD treatment while it decreased with Nec-1 treatment or *RIP3* deficiency (Figure 5). Thus, it is possible that to have effective long term neuroprotection autophagy and ROS may need to be modulated.

Surprisingly we observed that Nec-1 promotes axonal regeneration. Previous studies have shown that inhibition of caspase-6 or -8 stimulates a modest amount of axonal sprouting past the site of ON injury (20) and that inhibition of caspases-2 and -6 has a more substantial effect(22). In our studies, neither the pan-caspase inhibitor ZVAD nor the caspase-8-specific inhibitor z-IETD-FMK showed an effect over and above the vehicle. In contrast, Necrostatin exerted a more substantial effect on regeneration, which was minimally augmented by the addition of pan-caspase inhibitors. Surprisingly, although Nec-1 alone resulted in axonal regeneration, *Rip3*^*-/-*^ mice did not exhibit any. This could be due to the fact that Nec-1 is a RIP1 kinase inhibitor and thus effectors of RIP1 Kinase other than Rip3 may mediate the observed effect, or that other, as yet unknown targets of Nec-1 may be responsible for axonal rgenration.

The concept of neuroprotection, *i.e.*, the prevention of RGC death as a potential treatment for glaucoma patients who do not respond to current IOP lowering modalities or present with normal IOP, has become an extensive research field in glaucoma. Several neuroprotective drug-based monotherapy approaches have been proposed without showing any significant clinical efficacy (55–57). Our findings reveals the complexity and the redundancy of cell death pathways and suggests that multiple approaches may be required for effective RGC death prevention. Our study supports the idea that Nec-1 may act in a complementary fashion to other treatments that augment RGC survival and axon regeneration, yet even when identified apoptotic and necrotic pathways are blocked, RGC death continues. These results suggest the need to identify additional suppressors of cell death and stimulants of cell survival.

## EXPERIMENTAL PROCEDURES

### Animals

All animal experiments complied with the Association for Research in Vision and Ophthalmology for the use of animals in ophthalmic and vision research and were approved by the Animal Care and Use Committee of the Massachusetts Eye and Ear Infirmary (Boston, MA) and Boston Children’s Hospital (Boston, MA). C57BL/6 mice were purchased from Charles River (Wilmington, MA). *RIP3*^*-/-*^ mice on C57BL/6 background were provided from Dr. VM Dixit (Genentech, South San Francisco, CA) (58). The animals were fed standard laboratory chow and allowed free access to water in an air-conditioned room with a 12 h light/dark cycle. Except as noted otherwise, the animals were anesthetized with ketamine hydrochloride (30 mg/kg; Ketalar, Parke-Davis, Morris Plains, NJ) and xylazine hydrochloride (5 mg/kg; Rompun, Harver-Lockhart, Morris Plains, NJ) before all experimental manipulations.

### Mouse models of RGC loss

We utilized two animal models of RGC death. In the ON injury model, mice were anesthetized and subjected to severe crush injury at 1 mm distance from the eyeball for 15 sec using a cross-action forceps taking special care not to interfere with the blood supply (59). Injured mice were randomly divided into 4 groups for treatment: vehicle group (0.5% DMSO and 0.8% cyclodextrin in PBS, n = 6), ZVAD group (300 ⌠ M; Alexis, Plymouth Meeting, PA, n = 6), Nec-1 group (4 mM; a kind gift from Dr. J. Yuan, Harvard Medical School, Boston, MA, n = 6), and ZVAD plus Nec-1 group (n = 6). Soon after injury, each group received an intravitreal injection (2 ⌠ l) with the respective compounds..

### Quantification of RGC survival and measurement of IPL thickness

At indicated timepoints, eyes were enucleated and RGC loss was quantified from histological sections of mouse retina. Only transverse sections involving the optic disc were used for analysis and the fields corresponding to approximately 400 ⌠ m of both sides of retina extending from the ON head (2 points/section x 3 sections per eye, n = 6) were examined with an optical microscope (x40 objectives) (Figure S2). IPL thickness was measured with OpenLab software (Open Lab, Florence, Italy) (2 points/section x 3 sections per eye, n = 6). The ratio of IPL thickness was then calculated as a percentage of IPL thickness in the normal mouse eyes (n = 6).

### Quantitative real-time PCR

Total RNA extraction, cDNA synthesis and PCR amplification have been performed as previously reported (60). A real-time PCR assay was performed with Prism 7700 Sequence Detection System (Applied Biosystems, Foster City, CA). The primers are shown in Table S1. For relative comparison of each gene, we analyzed the Ct value of real-time PCR data with the ΔΔCt method normalizing by an endogenous control (®-actin).

### ELISA

The protein contents in retinal extract were determined with ELISA kits for protein carbonyls (Cell Biolabs, San Diego, CA) according to the manufacturer’s instructions.

### Western Blotting

Whole retinas were harvested and lysed for 30 min on ice in lysis buffer (50 mM Tris-HCl [pH 8], with 120 mM NaCl and 1% Nonidet P-40), supplemented with a mixture of proteinase inhibitors (Complete Mini; Roche Diagnostics, Basel, Switzerland). Thirty micrograms of protein per sample were separated in a 4-20% gradient sodium dodecyl sulfate-polyacrylamide gel (SDS-PAGE) (Invitrogen Corporation, Carlsbad, CA) electrophoresis and the proteins were electroblotted onto PVDF membranes. After 20 min incubation in blocking solution (Starting Block ™T20, Thermo Scientific, Waltham, MA), membranes were incubated with primary antibodies overnight at 4°C. Peroxidase-labeled secondary antibodies (Amersham Pharmacia Biotech, Piscataway, NJ) were used and proteins were visualized with enhanced chemiluminescence technique (Amersham Pharmacia Biotech). Full list of primary antibodies and working concentrations are shown in Table 1.

### Immunohistochemistry

Immunohistochemistry was performed as previously reported (60). After fixation and permeabilization, the sections were incubated with one of the primary antibodies (Table S1). An appropriate fluorophore-conjugated secondary antibody (Molecular Probes, Carlsbad, CA) was used to detect fluorescence using a confocal microscope (Leica Microsystems, Wetzler, Germany).

### Measurement of caspases-8, -9 and -3 activities

The activities of caspase-8, -9 and -3 were measured with the use of a commercially available kit according to the manufacturer’s instructions (APT171/131/139; Millipore, Billerica, MA). One day after injury, activities in the ON injured mice treated with vehicle, ZVAD, Nec-1 or ZVAD plus Nec-1 were normalized to their corresponding activities in non-ON injured retina (normal).

### TUNEL Analysis

TUNEL and quantification of TUNEL (+) cells were performed as previously described (61) by using the ApopTag Fluorescein *In Situ* Apoptosis Detection Kit (S7110; Chemicon International, Temecula, CA). The number of TUNEL (+) cells was calculated by the same way with the number of Brn3 (+) cells. A detailed description of the method is shown in Figure S2.

### TEM

TEM was performed as previously described (14). The eyes were enucleated and the posterior segments were fixed in 2.5% glutaraldehyde and 2% paraformaldehyde in 0.1 M cacodylate buffer with 0.08 M CaCl_2_ at 4°C. The eyes were post-fixed for 1.5 hours in 2% aqueous OsO_4_, dehydrated in ethanol and water, and embedded in EPON. Ultrathin sections were cut from blocks and stained with saturated, aqueous uranyl acetate and Sato’s lead stain. The specimens were observed with a Philips CM10 electron microscope. More than 50 RGCs per eye were photographed and subjected to quantification of cell death in a masked fashion. RGCs showing cellular shrinkage and nuclear condensation were defined as apoptotic cells, and RGCs associated with cellular and organelle swelling and discontinuities in nuclear and plasma membrane were defined as necrotic cells. Electron dense granular materials were labeled simply as end-stage unclassified cell death. Autophagosomes were defined as a double- or multi-membraned structure containing cytoplasmic material and/or organelles, and autolysosome was defined as cytoplasmic vesicle containing electron dense degraded material, as previously described (62).

### Long-term studies

Optic nerve crush surgery and intraocular injections were performed under general anesthesia. Adult mice are anesthetized with a combination of ketamine (1 mg/10 g body weight) and xylazine (0.1 mg/10 g body weight) given i.p. A conjunctival incision was made over the dorsal aspect of eye, which is then gently rotated downward in the orbit. The orbital muscles were separated to expose the optic nerve at its exit from the globe, which was then crushed for 5 seconds with jewelers’ forceps near the back of the eye (within 0.5 mm). Immediately. 3 days and 7 days after optic nerve crush, reagents were injected into the vitreous with a 30 1/2 gauge needle. Care was taken to avoid injuring lens and damaging the ophthalmic artery and retrobulbar sinus. 14 days after optic nerve injury, mice were sacrificed with an overdose of sodium pentobarbital and were perfused with saline and 4% paraformaldehyde (PFA). Optic nerves and eyes were dissected and postfixed in PFA. Nerves were impregnated with then 30% sucrose, embedded in OCT Tissue Tek Medium (Sakura Finetek), frozen, cut in the longitudinal plane at 14 μm, and mounted on coated slides. Regenerating axons were visualized by staining with a sheep antibody to GAP-43 followed by a fluorescently labeled secondary antibody. Axons were counted manually in at least eight longitudinal sections per case at prespecified distances from the injury site, and these numbers were converted into the number of regenerating axons at various distances (Leon et al., 2000).

The retinas were permeabilized in PBS 0.5% Triton by freezing them for 15 minutes at −80°C, rinsed in new PBS with 0.5% Triton, and incubated overnight at 4°C with goat antibody to Brn3a(C-20) (1: 100; Santa Cruz Biotechnology, inc.) and rabbit antibody to βIII-tubulin (1:500; Abcam) in blocking buffer (PBS, 2% bovine serum albumin, 2% Triton). Retinas were washed three times in PBS and incubated for 2 hours at room temperature with the Alexa Fluor-488-conjugated donkey anti-goat IgG antibody (1:500) and Alexa Fluor 594 donkey anti-rabbit (1:500) in blocking buffer. Finally, they were thoroughly washed in PBS and mounted vitreous side up on coated slides and covered with antifading solution. Images of eight prespecified areas, 2 mm from the optic disc were captured under fluorescent illumination (400X; E800; Nikon). Cells that stained positively for either Brn3a or βΠΙ -Tubulin or both were counted using NIH ImageJ software (Wayne Rasband, National Institutes of Health, Bethesda, MD). Cell densities were averaged across all eight areas, and data were represented as means ± SEM.

### Statistical Analysis

All values were expressed as the mean ± SD. Statistical differences between two groups were analyzed by Mann-Whitney *U* test. Multiple group comparison was performed by ANOVA followed by Tukey-Kramer adjustments. Differences were considered significant at *P* < 0.05.

## Acknowledgments

The authors thank Norman Michaud for technical assistance. This work was supported by Bacardi Fund (DV), Research to Prevent Blindness Foundation (DV), Lions Eye Research Fund (DV), Onassis Foundation (DV), Fight For Sight Grant in Aid (DV), Harvard Ophthalmology Department Support (DV), a Bausch & Lomb Vitreoretinal Fellowship (MK, Y. Morizane), The Department of Defense (Congressionally Directed Medical Research Program DM102446/Contract W81XWH-11-2-0023), the National Institutes of Health (MEEI Core Grant EY014104) and the Intellectual and Developmental Disabilities Research Center (IDDRC) of Boston Children’s Hospital (NIH P30 HD018655) for use of the Histology and Image Analysis Cores.

---

**Suppl Fig1.**
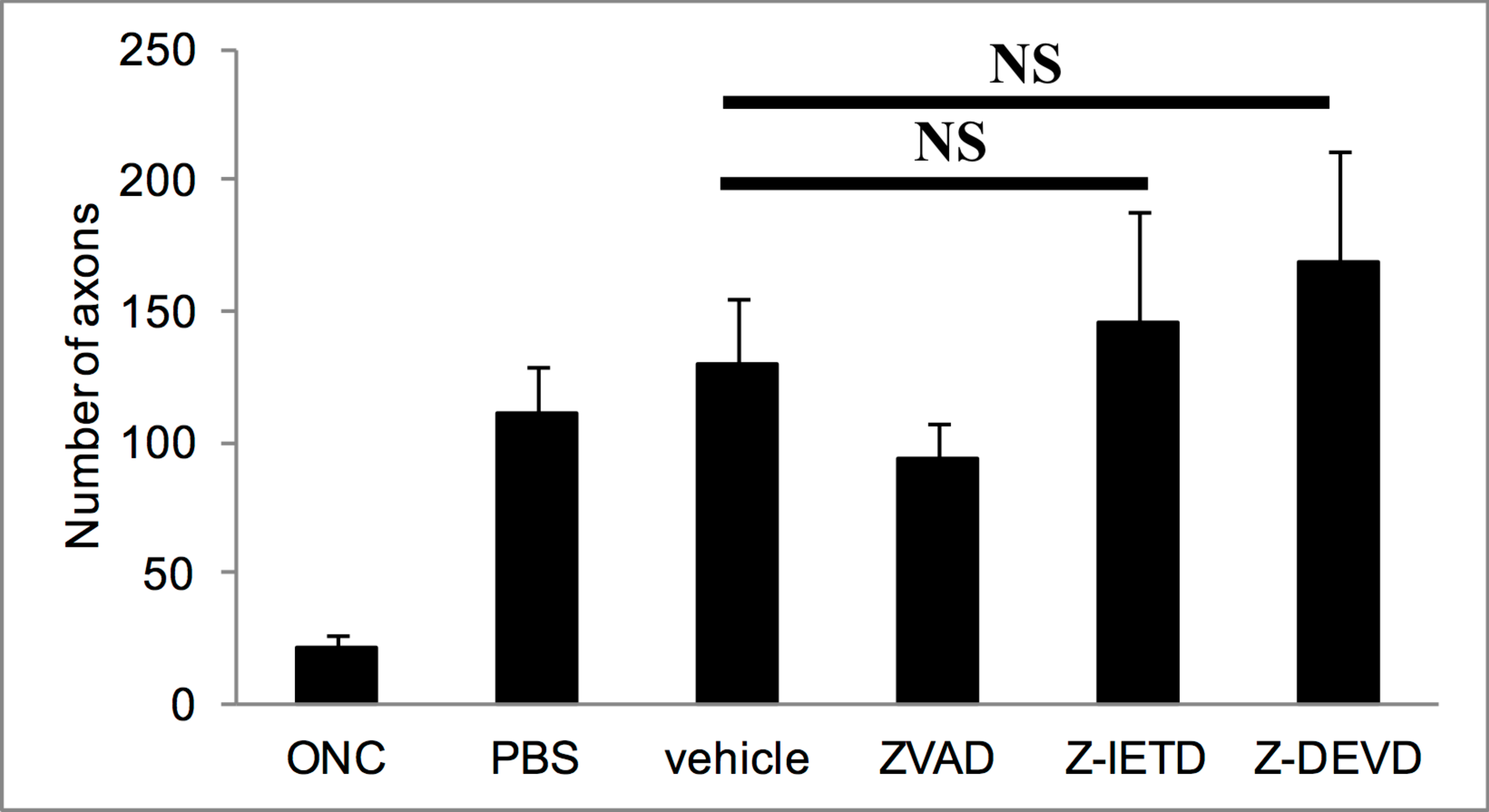

**supplementary FIGURE2.**
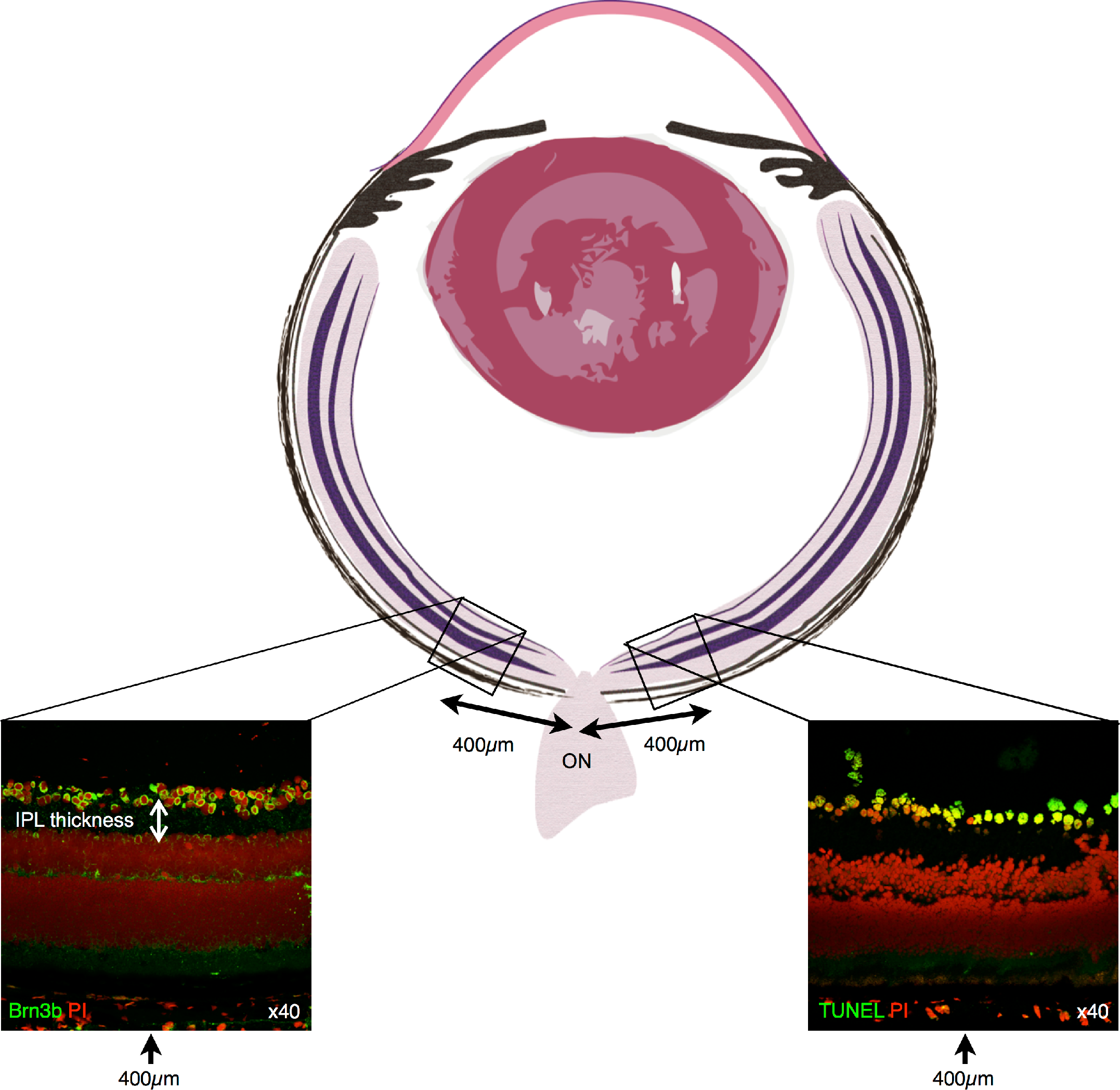

## REFERENCES

1. Resnikoff S, et al (2004) Global data on visual impairment in the year 2002. Bull World Health Organ 82(11):844–851.

2. Quigley HA & Broman AT (2006) The number of people with glaucoma worldwide in 2010 and 2020. Br J Ophthalmol 90(3):262–267.

3. Weinreb RN & Khaw PT (2004) Primary open-angle glaucoma. Lancet 363 (9422): 1711–1720.

4. Kwon YH, Fingert JH, Kuehn MH, & Alward WL (2009) Primary open-angle glaucoma. N Engl J Med 360(11):1113–1124.

5. Kroemer G, et al (2009) Classification of cell death: recommendations of the Nomenclature Committee on Cell Death 2009. Cell Death Differ 16(1):3–11.

6. Degterev A & Yuan J (2008) Expansion and evolution of cell death programmes. Nat Rev Mol Cell Biol 9(5):378–390.

7. Holler N, et al (2000) Fas triggers an alternative, caspase-8-independent cell death pathway using the kinase RIP as effector molecule. Nat Immunol 1(6):489–495.

8. Levine B & Kroemer G (2008) Autophagy in the pathogenesis of disease. Cell 132(1):27–42.

9. Tezel G & Wax MB (1999) Inhibition of caspase activity in retinal cell apoptosis induced by various stimuli in vitro. Invest Ophthalmol Vis Sci 40(ll):2660–2667.

10. Vandenabeele P, Declercq W, Van Herreweghe F, & Vanden Berghe T (2010) The role of the kinases RIP1 and RIP3 in TNF-induced necrosis. Sci Signal 3(115):re4.

11. Cho YS, et al. (2009) Phosphorylation-driven assembly of the RIP1-RIP3 complex regulates programmed necrosis and virus-induced inflammation. Cell 137(6):1112–1123.

12. He S, et al (2009) Receptor interacting protein kinase-3 determines cellular necrotic response to TNF-alpha. Cell 137(6):1100–1111.

13. Zhang DW, et al (2009) RIP3, an energy metabolism regulator that switches TNF-induced cell death from apoptosis to necrosis. Science 325(5938):332–336.

14. Trichonas G, et al (2010) Receptor interacting protein kinases mediate retinal detachment-induced photoreceptor necrosis and compensate for inhibition of apoptosis. Proc Natl Acad Sci USA.

15. Degterev A, et al (2005) Chemical inhibitor of nonapoptotic cell death with therapeutic potential for ischemic brain injury. Nat Chem Biol 1(2):112–119.

16. Nakazawa T, et al (2006) Tumor necrosis factor-alpha mediates oligodendrocyte death and delayed retinal ganglion cell loss in a mouse model of glaucoma. Neurosci 26(49):12633–12641.

17. Roh M, et al (2012) Etanercept, a widely used inhibitor of tumor necrosis factor-alpha (TNF-alpha), prevents retinal ganglion cell loss in a rat model of glaucoma. PLoS One 7(7):e40065.

18. Tezel G, Li LY, Patil RV, & Wax MB (2001) TNF-alpha and TNF-alpha receptor-1 in the retina of normal and glaucomatous eyes. Invest Ophthalmol Vis Sci 42(8):1787–1794.

19. Tezel G (2008) TNF-alpha signaling in glaucomatous neurodegeneration. Prog Brain Res 173:409–421.

20. Monnier PP, et al. (2011) Involvement of caspase-6 and caspase-8 in neuronal apoptosis and the regenerative failure of injured retinal ganglion cells. JNeurosci 31(29):10494–10505.

21. Ahmed Z, et al. (2011) Ocular neuroprotection by siRNA targeting caspase-2. Cell Death Dis 2:e173.

22. Vigneswara V, et al. (2014) Combined suppression of CASP2 and CASP6 protects retinal ganglion cells from apoptosis and promotes axon regeneration through CNTF-mediated JAK/STAT signalling. Brain 137(Pt 6):1656–1675.

23. Nickells RW (1999) Apoptosis of retinal ganglion cells in glaucoma: an update of the molecular pathways involved in cell death. Surv Ophthalmol 43 Suppl 1:S151–161.

24. Kermer P, et al. (1999) Activation of caspase-3 in axotomized rat retinal ganglion cells in vivo. FEBS Lett 453(3):361–364.

25. Chaudhary P, Ahmed F, Quebada P, & Sharma SC (1999) Caspase inhibitors block the retinal ganglion cell death following optic nerve transection. Brain Res Mol Brain Res 67(1):36–45.

26. Tezel G & Yang X (2004) Caspase-independent component of retinal ganglion cell death, in vitro. Invest Ophthalmol Vis Sci 45(ll):4049–4059.

27. Libby RT, et al. (2005) Susceptibility to neurodegeneration in a glaucoma is modified by Bax gene dosage. PLoS Genet 1(1):17–26.

28. Watkins TA, et al. (2013) DLK initiates a transcriptional program that couples apoptotic and regenerative responses to axonal injury. Proc Natl AcadSciUSA 110(10):4039–4044.

29. Welsbie DS, et al. (2013) Functional genomic screening identifies dual leucine zipper kinase as a key mediator of retinal ganglion cell death. Proc Natl Acad Sci USA 110(10):4045–4050.

30. Quigley HA, et al. (1995) Retinal ganglion cell death in experimental glaucoma and after axotomy occurs by apoptosis. Invest Ophthalmol Vis Sci 36(5):774–786.

31. Grasl-Kraupp B, et al. (1995) In situ detection of fragmented DNA (TUNEL assay) fails to discriminate among apoptosis, necrosis, and autolytic cell death: a cautionary note. Hepatology 21(5):1465–1468.

32. Unal Cevik I & Dalkara T (2003) Intravenously administered propidium iodide labels necrotic cells in the intact mouse brain after injury. Cell Death Differ 10(8):928–929.

33. Kelliher MA, et al. (1998) The death domain kinase RIP mediates the TNF-induced NF-kappaB signal. Immunity 8(3):297–303.

34. Kim YS, Morgan MJ, Choksi S, & Liu ZG (2007) TNF-induced activation of the Nox1 NADPH oxidase and its role in the induction of necrotic cell death. Mol Cell 26(5):675–687.

35. Li Y, Irwin N, Yin Y, Lanser Μ, & Benowitz LI (2003) Axon regeneration in goldfish and rat retinal ganglion cells: differential responsiveness to carbohydrates and cAMP. J Neurosci 23(21):7830–7838.

36. Yin Y, et al. (2009) Oncomodulin links inflammation to optic nerve regeneration. Proc Natl AcadSci USA 106(46):19587–19592.

37. Kurimoto T, et al. (2013) Neutrophils express oncomodulin and promote optic nerve regeneration. Neurosci 33(37):14816–14824.

38. Quigley HA, Addicks EM, Green WR, & Maumenee AE (1981) Optic nerve damage in human glaucoma. II. The site of injury and susceptibility to damage. Arch Ophthalmol 99(4):635–649.

39. Hernandez MR (1993) Extracellular Matrix Macromolecules of the Lamina Cribrosa: A Pressure-sensitive Connective Tissue. Glaucoma 2(l):50–57.

40. Sawada H, Fukuchi T, Tanaka T, & Abe H (2010) Tumor necrosis factor-alpha concentrations in the aqueous humor of patients with glaucoma. Invest Ophthalmol Vis Sci 51(2):903–906.

41. Funayama T, et al. (2004) Variants in optineurin gene and their association with tumor necrosis factor-alpha polymorphisms in Japanese patients with glaucoma. Invest Ophthalmol Vis Sci 45(12):4359–4367.

42. Yuan L & Neufeld AH (2000) Tumor necrosis factor-alpha: a potentially neurodestructive cytokine produced by glia in the human glaucomatous optic nerve head. Glia 32(1):42–50.

43. Yuan L & Neufeld AH (2001) Activated microglia in the human glaucomatous optic nerve head? Neurosci Res 64(5):523–532.

44. Ju KR, Kim HS, Kim JH, Lee NY, & Park CK (2006) Retinal glial cell responses and Fas/FasL activation in rats with chronic ocular hypertension. Brain Res 1122(1):209–221.

45. Wax MB, et al. (2008) Induced autoimmunity to heat shock proteins elicits glaucomatous loss of retinal ganglion cell neurons via activated T-cell-derived fas-ligand. J Neurosci 28(46):12085–12096.

46. Jiang B, et al. (2010) Neuroinflammation in advanced canine glaucoma. Mol Vis 16:2092–2108.

47. Gregory MS, et al. (2011) Opposing roles for membrane bound and soluble Fas ligand in glaucoma-associated retinal ganglion cell death. PLoS One 6(3):e17659.

48. Yang X, et al. (2011) Neurodegenerative and inflammatory pathway components linked to TNF-alpha/TNFR1 signaling in the glaucomatous human retina. Invest Ophthalmol Vis Sci 52(ll):8442–8454.

49. Murakami Y, et al. (2012) Receptor interacting protein kinase mediates necrotic cone but not rod cell death in a mouse model of inherited degeneration. Proc Natl Acad Sci USA 109(36):14598–14603.

50. Tezel G & Yang X (2005) Comparative gene array analysis of TNF-alpha-induced MAPK and NF-kappaB signaling pathways between retinal ganglion cells and glial cells. Exp Eye Res 81(2):207–217.

51. Degterev A, et al. (2008) Identification of RIP1 kinase as a specific cellular target of necrostatins. Nat Chem Biol 4(5):313–321.

52. Joo CK, et al (1999) Necrosis and apoptosis after retinal ischemia: involvement of NMDA-mediated excitotoxicity and p53. Invest Ophthalmol Vis Sci 40(3):713–720.

53. Saggu SK, Chotaliya HP, Blumbergs PC, & Casson RJ (2010) Wallerian-like axonal degeneration in the optic nerve after excitotoxic retinal insult: an ultrastructural study. BMC Neurosci 11:97.

54. Yu L, et al (2006) Autophagic programmed cell death by selective catalase degradation. Proc Natl Acad Sci USA 103(13):4952–4957.

55. Guo L, et al (2006) Assessment of neuroprotective effects of glutamate modulation on glaucoma-related retinal ganglion cell apoptosis in vivo. Invest Ophthalmol Vis Sci 47(2):626–633.

56. Osborne NN (2009) Recent clinical findings with memantine should not mean that the idea of neuroprotection in glaucoma is abandoned. Acta Ophthalmol 87(4):450–454.

57. Saylor M, McLoon LK, Harrison AR, & Lee MS (2009) Experimental and clinical evidence for brimonidine as an optic nerve and retinal neuroprotective agent: an evidence-based review. Arch Ophthalmol 127(4):402–406.

58. Newton K, Sun X, & Dixit VM (2004) Kinase RIP3 is dispensable for normal NF-kappa Bs, signaling by the B-cell and T-cell receptors, tumor necrosis factor receptor 1, and Toll-like receptors 2 and 4. Mol Cell Biol 24(4):1464–1469.

59. Levkovitch-Verbin H (2004) Animal models of optic nerve diseases. Eye (Lond) 18(11):1066–1074.

60. Kayama M, et al (2010) Transfection with pax6 gene of mouse embryonic stem cells and subsequent cell cloning induced retinal neuron progenitors, including retinal ganglion cell-like cells, in vitro. Ophthalmic Res 43(2):79–91.

61. Nakazawa T, et al (2007) Monocyte chemoattractant protein 1 mediates retinal detachment-induced photoreceptor apoptosis. Proc Natl Acad Sci U S A 104(7):2425–2430.

62. Eskelinen EL (2008) To be or not to be? Examples of incorrect identification of autophagic compartments in conventional transmission electron microscopy of mammalian cells. Autophagy 4(2):257–260.

